# A combination of transcription factors mediates inducible interchromosomal pairing

**DOI:** 10.1101/385047

**Authors:** Seungsoo Kim, Maitreya J Dunham, Jay Shendure

## Abstract

Remodeling of the three-dimensional organization of a genome has been previously described (*e.g.* condition-specific pairing or looping), but it remains unknown which factors specify and mediate such shifts in chromosome conformation. Here we describe an assay, MAP-C (Mutation Analysis in Pools by Chromosome conformation capture), that enables the simultaneous characterization of hundreds of *cis* or *trans*-acting mutations for their effects on a chromosomal contact or loop. As a proof of concept, we applied MAP-C to systematically dissect the molecular mechanism of inducible interchromosomal pairing between *HAS1pr-TDA1pr* alleles in *Saccharomyces* yeast. We identified three transcription factors, Leu3, Sdd4 (Ypr022c), and Rgt1, whose collective binding to nearby DNA sequences is necessary and sufficient for inducible pairing between binding site clusters. Rgt1 contributes to the regulation of pairing, both through changes in expression level and through its interactions with the Tup1/Ssn6 repressor complex. *HAS1pr-TDA1pr* is the only locus with a cluster of binding site motifs for all three factors in both *S. cerevisiae* and *S. uvarum* genomes, but the promoter for *HXT3*, which contains Leu3 and Rgt1 motifs, also exhibits inducible homolog pairing. Altogether, our results demonstrate that specific combinations of transcription factors can mediate condition-specific interchromosomal contacts, and reveal a molecular mechanism for interchromosomal contacts and mitotic homolog pairing.

## Introduction

The three-dimensional organization of the genome within the nucleus is non-random (Bonev and Cavalli, 2016). Although many features of this conformation are largely conserved across cell types and conditions (Rao et al., 2014; Schmitt et al., 2016a), some chromatin loops and contacts form specifically in response to signals such as differentiation (Bonev et al., 2017; Schmitt et al., 2016a; Stadhouders et al., 2018), changes in nutrient availability (Brickner et al., 2015, 2016), heat shock (Chowdhary et al., 2017), drugs (D’Ippolito et al., 2018), meiosis (Muller et al., 2018), or circadian rhythms (Kim et al., 2018). This dynamic three-dimensional organization of the genome plays a role in regulating gene expression in diverse organisms. In multicellular organisms, active developmental gene promoters form long-range loops with specific enhancer elements (Bonev et al., 2017), and this looping is in some cases sufficient for activation (Deng et al., 2014). In the budding yeast *Saccharomyces cerevisiae*, a well-studied model of genome conformation, genes targeted to nuclear pores are activated (Taddei et al., 2006), whereas those at the nuclear periphery are repressed (Andrulis et al., 1998). However, the mechanisms driving such conformational reorganizations remain largely unknown.

Transcription factors (TFs) are attractive candidates for orchestrating dynamic changes in chromatin conformation, given their site-specific DNA binding and changes in abundance or activity in response to differentiation and cellular signals (Lambert et al., 2018). For many conditions, it remains unknown exactly which TFs bind to any given locus; although binding site motifs are known for many TFs, a motif match is insufficient to fully predict TF binding (Le et al., 2018; Levo et al., 2015; Slattery et al., 2014). Even if the set of TFs bound to each locus is known, it is unclear which TFs are capable of forming chromosomal contacts. Furthermore, DNA-bound TFs can recruit other cofactor proteins that can mediate chromosomal contacts (Deng et al., 2014), but our understanding of TF-cofactor interactions remains incomplete.

Among inducible chromosomal contacts and loops, interchromosomal contacts are particularly poorly understood. Although interchromosomal contacts, such as those between actively transcribed genes or homolog pairing in mitotically dividing yeast, have been observed by microscopy (Burgess et al., 1999; Lim et al., 2018; Maass et al., 2018), many are not detectable using 3C (chromosome conformation capture) technologies. Some have argued that these contacts might be a result of fluorescent in situ hybridization (FISH) artifacts (Lorenz et al., 2003) or the insertion of binding site arrays often used for live-cell imaging (Mirkin et al., 2014). A more detailed understanding of the molecular components of interchromosomal contacts is needed to resolve this ambiguity.

We recently identified a novel example of inducible interchromosomal pairing between homologous copies of the *HAS1pr-TDA1pr* locus in diploid *Saccharomyces* yeasts (Kim et al., 2017). This interaction occurs in saturated culture conditions, requires the 1 kb intergenic region between the *HAS1* and *TDA1* coding sequences, and is detectable by both Hi-C and microscopy.

The condition-specificity and dependence on intergenic sequence led us to hypothesize that one or more TFs might mediate this pairing. Although yeast TF binding is well-characterized for standard growth conditions (Badis et al., 2008), TF binding has not been systematically experimentally measured in saturated culture conditions. In fact, within the 1 kb region required for *HAS1pr-TDA1pr* pairing, dozens of TFs have at least one motif match. Furthermore, even if the TFs bound to this region were known, it would remain unclear which subset played a role in mediating inducible interchromosomal pairing.

Here we describe a method that enables one to simultaneously test hundreds of *cis* or *trans*-acting mutations for their effects on a chromosomal contact or loop of interest. As a proof of concept, we applied this method, which we call Mutation Analysis in Pools by Chromosome conformation capture (MAP-C), to characterize the molecular components mediating *HAS1pr-TDA1pr* pairing. Through saturating mutagenesis of the regulatory region that mediates the interchromosomal pairing (*cis* MAP-C), we identify sequence motifs potentially corresponding to TFs. By also testing the effects of knocking out these candidate TFs (*trans* MAP-C), we confirm that binding of three TFs—Leu3, Sdd4 (Ypr022c), and Rgt1—are individually necessary and collectively sufficient for inducible interchromosomal pairing. We further use *trans* MAP-C to interrogate how known interaction partners of Rgt1 regulate pairing. Searching for additional loci with heterotypic clusters of binding sites for these TFs, we identify *HXT3pr* as another locus exhibiting inducible pairing. Together, our results demonstrate how a combination of TFs can mediate inducible interchromosomal pairing and the utility of a pooled mutant approach to studying chromosome conformation.

## Results

### A pooled approach to systematically dissect chromosome conformation

In order to identify and dissect the molecular mechanisms underlying chromosome conformation, experiments involving perturbations (*e.g.* mutations) are needed. However, despite the advances in chromosome conformation capture (3C) technology over the last two decades (Schmitt et al., 2016b; de Wit and de Laat, 2012), each experiment remains limited to one sample, limiting the testing of perturbations to top candidate genes or elements (Nora et al., 2017; Schwarzer et al., 2017; Weintraub et al., 2017).

To address this limitation and enable systematic screens, we developed MAP-C, an assay in which hundreds of mutations are simultaneously tested for their effects on a single chromosomal contact of interest (**Figure 1A**). The *cis* version of MAP-C, which we describe first, allows for the generation of an allelic series in a cost-effective, pooled manner (*e.g.* by array-synthesized oligonucleotide pools or error-prone PCR) or individually, as long as the variants correspond to one of the regions involved in the chromosomal contact. The resulting mutant pool is then subjected to the 3C assay, and the region containing the genetic variants is amplified using two different primer pairs: the first (3C library) amplifies a specific ligation product, and the second (genomic library) amplifies regardless of ligation. These amplification products are deeply sequenced to measure the abundance of each variant in the 3C library, which is normalized to its abundance in the genomic library. The relative extent to which sequence variants participate in the chromosomal contact of interest is proportional to their normalized representation in the 3C library.

**Figure 1.**
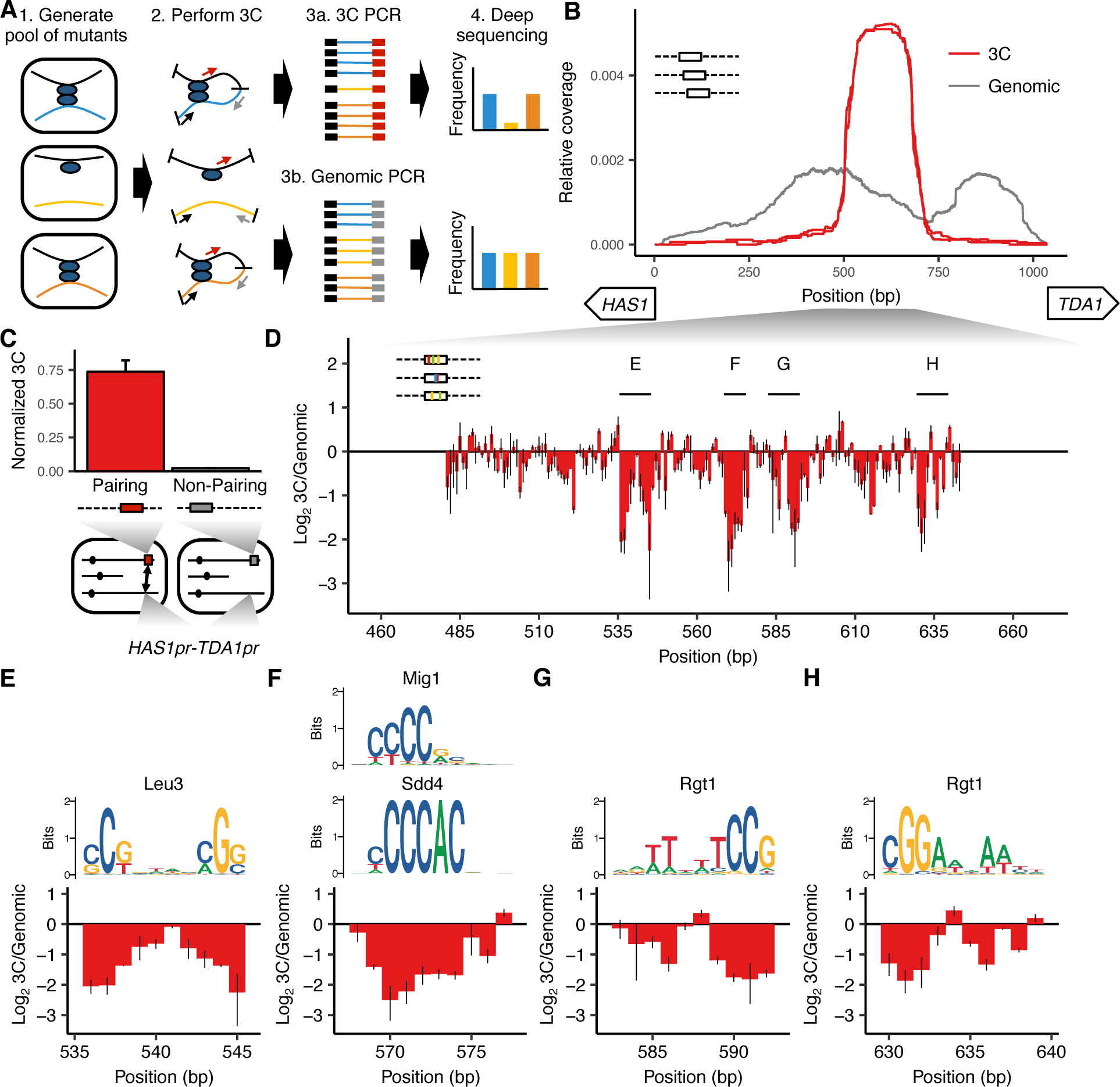
**MAP-C identifies DNA sequences necessary and sufficient for inducible pairing between *HAS1pr-TDA1pr* alleles.** (A) Schematic of *cis* flavor of MAP-C method. Colored lines indicate mutant DNA sequences, and thin arrows indicate primers. (B) 178 bp subsequences from the *S. cerevisiae HAS1pr-TDA1pr* region sufficient for pairing with the *S. uvarum HAS1pr-TDA1pr*, shown as read coverage of the 3C (red) and genomic (gray) libraries, normalized to sum to 1. The two lines for each color represent technical replicates. (C) 3C for contact between *HAS1pr-TDA1pr* and a pairing (red) or non-pairing (gray) sequence (coordinates shown in **Figure S1C**) integrated at the *FIT1* locus in haploid *S. cerevisiae*, normalized to contacts between *FIT1* and *HLR1* 10 kb away. Bars indicate mean ± s.d. of technical triplicates. (D) Base-pairs necessary for pairing, shown as ratio of the total substitution frequency at each position in the 3C library compared to the genomic library. Error bars indicate the two technical replicates. Start positions and orientations of *HAS1* and *TDA1* coding sequences are shown on x-axis. (E-H) Selected regions from panel D are highlighted, with sequence logos for matching transcription factor motifs.

### A cluster of TF motifs is necessary and sufficient for *HAS1pr-TDA1pr* pairing

As a first test of *cis* MAP-C, we sought to systematically dissect the conserved pairing between *HAS1pr-TDA1pr* homologs in diploid *Saccharomyces* yeasts grown to saturation. We recently used Hi-C of *S. cerevisiae* x *S. uvarum* hybrids to discover this homolog pairing interaction, and used furthermore identified a 1,038 bp noncoding region that was necessary and sufficient for pairing (Kim et al., 2017).

To find a minimal subsequence of the 1,038 bp *HAS1pr-TDA1pr* region that is sufficient to pair with other *HAS1pr-TDA1pr* alleles, we replaced the native *S. cerevisiae HAS1pr-TDA1pr* locus with a library containing each of 861 tiling 178 bp subsequences of the 1,038 bp region (along with a G418 resistance cassette and restriction site), in *S. cerevisiae* x *S. uvarum* hybrid yeast. We then performed *cis* MAP-C for pairing of the modified locus with the *S. uvarum* copy of *HAS1pr-TDA1pr* on a saturated culture of the pool, in two replicates (**Figure 1B**). Compared to the genomic libraries, the 3C libraries were highly enriched for a narrow region spanning ~500 to ~700 bp from the *HAS1* coding sequence, with a plateau between ~525 to ~675 bp, consistent with that region being the only subsequence shorter than 178 bp sufficient for pairing (termed the “minimal pairing region” below). To confirm that this pattern of enrichment is specific to *HAS1pr-TDA1pr* homolog pairing rather than underlying all of its chromosomal contacts, we repeated the assay with a pair of primers amplifying an intrachromosomal contact (**Figure S1A and B**). In this “off-target” control, normalized coverage from the 3C library was uniform, suggesting that most variants are capable of intrachromosomal looping, but only those containing minimal pairing region are capable of interchromosomal pairing with the other *HAS1pr-TDA1pr* allele.

Since our initial experiment was performed in the endogenous genomic context, other DNA sequences outside but near the *HAS1pr-TDA1pr* locus could be required in addition to the minimal pairing region. We therefore tested whether inserting a 184 bp sequence that included the minimal pairing region into an ectopic location, the gene *FIT1* (*YDR534C*), would induce pairing with the native *HAS1pr-TDA1pr* locus in saturated cultures of haploid *S. cerevisiae* (**Figure S1C**). As a negative control, we inserted an equivalently sized subsequence insufficient for pairing into the same locus (**Figure S1C**). Indeed, insertion of the minimal pairing region led to a >30-fold increase in 3C signal for pairing with *HAS1pr-TDA1pr* as compared to the negative control (**Figure 1C**).

We next sought to obtain a base-pair resolution map of the DNA sequences necessary for pairing. We used error-prone PCR to generate variants of a 207 bp region containing the minimal pairing region, with an average of 1.49 substitutions (range 0-14) per template (**Figure S1D**). We inserted this variant library in place of the native *S. cerevisiae HAS1pr-TDA1pr* sequence as before, and performed *cis* MAP-C. Plotting the ratio of total substitution abundance in the 3C and genomic libraries at each mutagenized position identified five clusters of positions showing strong depletion of substitutions in the 3C libraries, indicating that they are required for *HAS1pr-TDA1pr* pairing (**Figure 1D**). All five clusters corresponded to one or more TF motifs: the first two clusters together aligned to a Leu3 motif (**Figure 1E**), the third to several similar motifs, including Sdd4 (Ypr022c) and Mig1 (**Figure 1F**), and the last two to Rgt1 motifs in opposite orientations (**Figure 1G and H**). None of these mutations had an effect on intrachromosomal looping (**Figure S1E**), and all of the clusters were reproduced using an alternative mutagenesis strategy (programmed 3 bp substitutions) (**Figure S2**). Interestingly, a third Rgt1 motif and a second Sdd4/Mig1 motif mutagenized only in our validation experiment were not required for pairing, suggesting that either not all motifs in this region are bound by the same TFs or not all bound TFs are involved in mediating homolog pairing at this locus.

Thus, using *cis* MAP-C, we identified a ~150 bp subsequence of *HAS1pr-TDA1pr* sufficient for pairing, containing four required TF motif occurrences. If these TF motifs are together sufficient for pairing, we would expect that 1) they are only observed in a cluster in the minimal pairing region and not elsewhere in the *HAS1pr-TDA1pr*, and 2) they are present in a cluster in the *S. uvarum* copy of this region, and potentially other *Saccharomyces* as well. Indeed, the motifs are isolated to the central region of *HAS1pr-TDA1pr* in *S. cerevisiae*, *S. uvarum*, and in fact across all *Saccharomyces* species (**Figure S3**).

### Three transcription factors are required for pairing

Despite identifying the TF motifs required for *HAS1pr-TDA1pr* homolog pairing at base-pair resolution with *cis* MAP-C, the the redundancy among TF motifs made it difficult to definitively identify the TFs involved. To address this, we developed a modified version of MAP-C that could assay *trans* knockouts of genes spread across the genome for their effects on a specific chromosomal contact (*trans* MAP-C). With *trans* MAP-C, each gene knockout is uniquely associated with a short barcode sequence near a copy of the pairing sequence, which is then assayed by 3C for interactions with a chromosomal contact of interest (**Figure 2A**). Mutations that affect pairing frequency modulate the abundance of their corresponding barcodes in the 3C products, which can be readily quantified by deep sequencing.

**Figure 2.**
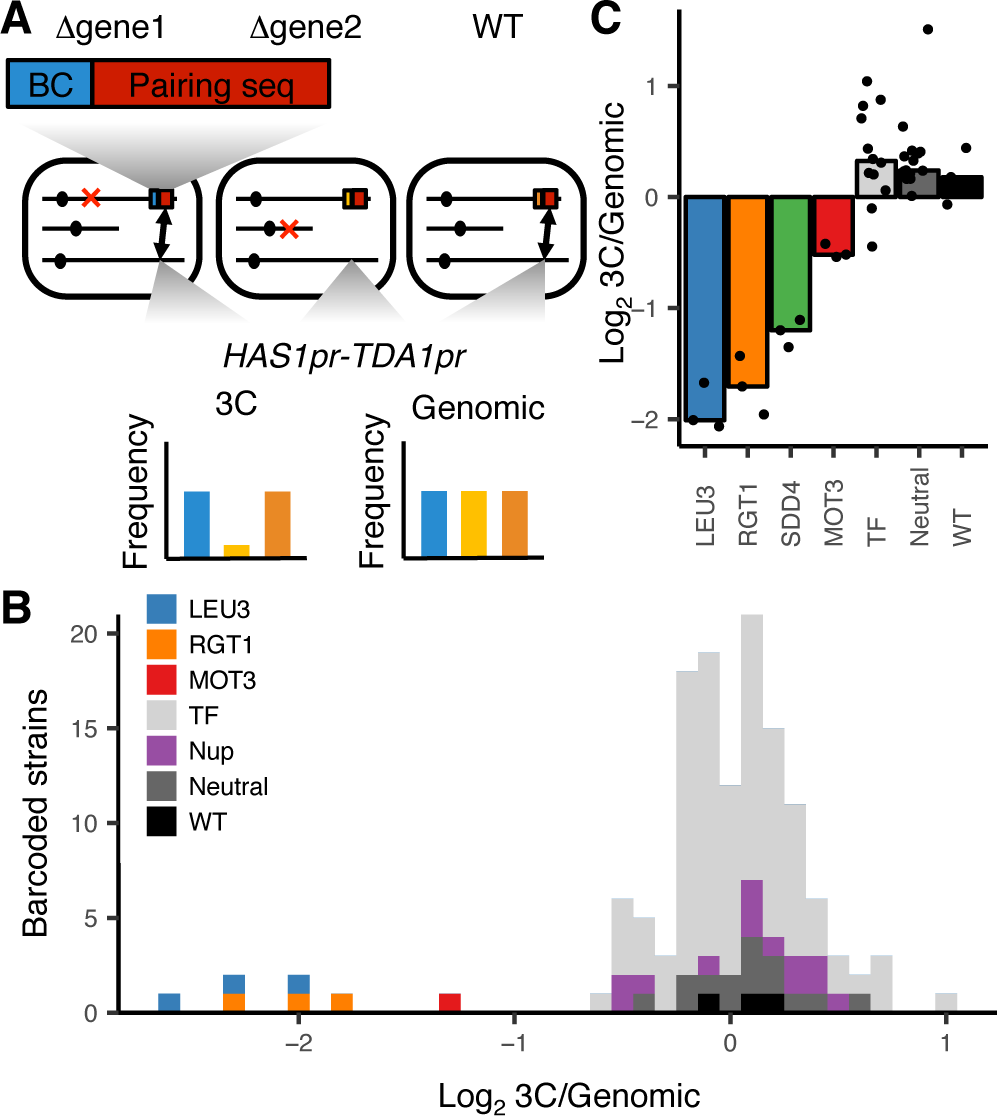
***Trans* factors required for *HAS1pr-TDA1pr* pairing.** (A) Schematic of barcoded knockout in *trans* flavor of MAP-C. Red Xs indicate gene knockouts; red boxes indicate pairing sequence (**Figure S1C**); BC indicates barcode; double-headed arrows indicate the presence of chromosomal contacts. (B) Histogram of relative abundance of each barcoded gene knockout strain in 3C library compared to the genomic library, excluding strains below a frequency of 0.3% of the pool. Barcode replicates are shown as separate squares in histogram. *LEU3*, *RGT1*, and *MOT3* are highlighted individually; TF indicates other transcription factors; Nup indicates nuclear pore complex components; Neutral indicates fitness-neutral negative controls (**Figure S5A**). (C) Validation TF knockout screen, shown as bar plot of median relative abundance in 3C library compared to the genomic library, with overlaid scatter plot of individual barcoded strains. TF includes *MIG1*, *VHR1*, *CBF1*, and *YGR067C*.

We first tested this approach by inserting the minimal pairing region (**Figure S1B**) into the common KanMX drug resistance cassette in a pilot subset of 10 TF knockout strains from the haploid yeast deletion collection (Giaever et al., 2002; Mirkin et al., 2013). We then assayed these constructs for interactions with the native *HAS1pr-TDA1pr* region, using the existing gene-specific barcodes to measure strain abundances in each library. In this approach, the frequency of pairing is confounded by the different genomic location of the pairing sequence in each strain. Therefore, for each of the 10 TFs we targeted, we included as controls up to 6 neighboring genes, which should have a similar genomic location effect as the targeted gene (**Figure S4A**). We hypothesized that due to the Rabl orientation of yeast chromosomes, in which centromeres are clustered together (Duan et al., 2010), centromere-proximal regions would interact less with *HAS1pr-TDA1pr*, which is centromere-distal. Indeed, the most centromere-distal gene knockouts interacted ~4-fold more with the *HAS1pr-TDA1pr* locus than the most centromere-proximal gene knockouts (**Figure S4B**). Of the 7 TF gene knockouts that were measured in our assay (3 dropped out during library construction, including *LEU3*), six had no substantial difference in pairing strength compared to their genomic neighbors (**Figure S4C**). However, knockout of *RGT1* led to a ~20-fold decrease in pairing strength, consistent with Rgt1 binding its cognate motifs in the minimal pairing region (**Figure 2G and H**).

Next, we expanded our *trans* MAP-C screen to include the majority of known nonessential TFs with motifs (de Boer and Hughes, 2012). To avoid the confounding effect of genomic location, we inserted a barcoded pairing sequence construct into a fixed locus and associated each of these barcodes with the cognate knockout by individually transforming each knockout strain with a unique barcode. We tested a total of 109 TF gene knockouts, as well as 15 nuclear pore complex components, 8 fitness neutral negative controls, and a wild-type control, with multiple barcode replicates for controls and expected hits (**Figure S5A**). As expected, most barcoded strains were equally abundant in the 3C and genomic libraries, indicating that the corresponding knockout did not impact *HAS1pr-TDA1pr* pairing. However, *LEU3* and *RGT1* knockouts were depleted ~4- fold from the 3C libraries, suggesting that they are required for pairing (**Figure 2B**). In addition, deletion of *MOT3* modestly decreased pairing (~2.5-fold). Two other knockouts, *VHR1* and *CBF1*, appeared to also decrease pairing, but were at low abundances and might reflect noise (**Figure S5B**).

Surprisingly, none of the TFs required for pairing had a motif matching the sequence CCCCAC (**Figures 1F, 2B, and S4C**). However, two putative TFs, *YPR022C* (*SDD4*) and *YGR067C*, with high scoring motif matches were excluded in the initial screens due to their lack of annotations. Therefore, we repeated our fixed-locus TF knockout screen with a limited set of genes, including the two putative TFs and additional replicates for the hits *MOT3*, *VHR1*, and *CBF1*. We found that indeed, *SDD4* is required for pairing, suggesting that it is the *trans* acting factor that binds the CCCCAC motif (**Figure 2C**). *MOT3* once again exhibited modest depletion, suggesting a minor, perhaps indirect role in pairing, whereas *VHR1* and *CBF1* displayed no depletion (**Figure 2C**).

Thus, deletion of either the genes encoding Leu3, Rgt1, and Sdd4, or mutations in their binding sites located within a ~150 bp region of the *HAS1pr-TDA1pr* locus, disrupt inducible homolog pairing between *HAS1pr-TDA1pr* alleles. Our results are consistent with these three TFs mediating chromosomal contacts by binding to DNA and interacting directly with one another and/or indirectly through cofactors.

### Changes in Rgt1 abundance and recruitment of Tup1/Ssn6 regulate pairing

We next sought to explore the mechanisms that regulate pairing. We hypothesized that the TF expression levels might regulate the strength of *HAS1pr-TDA1pr* pairing, paralleling the regulation of TF-mediated transcriptional activation. To test this hypothesis, we analyzed RNA-seq data for haploid *S. cerevisiae* in pairing and non-pairing conditions (saturated and exponentially growing cultures, respectively) (Kim et al., 2017). *RGT1* and *SDD4* were upregulated in saturated cultures, ~2-fold and ~6-fold, respectively, whereas *LEU3* transcript levels remained constant (**Figure 3A**). These results are consistent with the hypothesis that increased transcription of TFs, particularly *RGT1*, regulates pairing strength.

**Figure 3.**
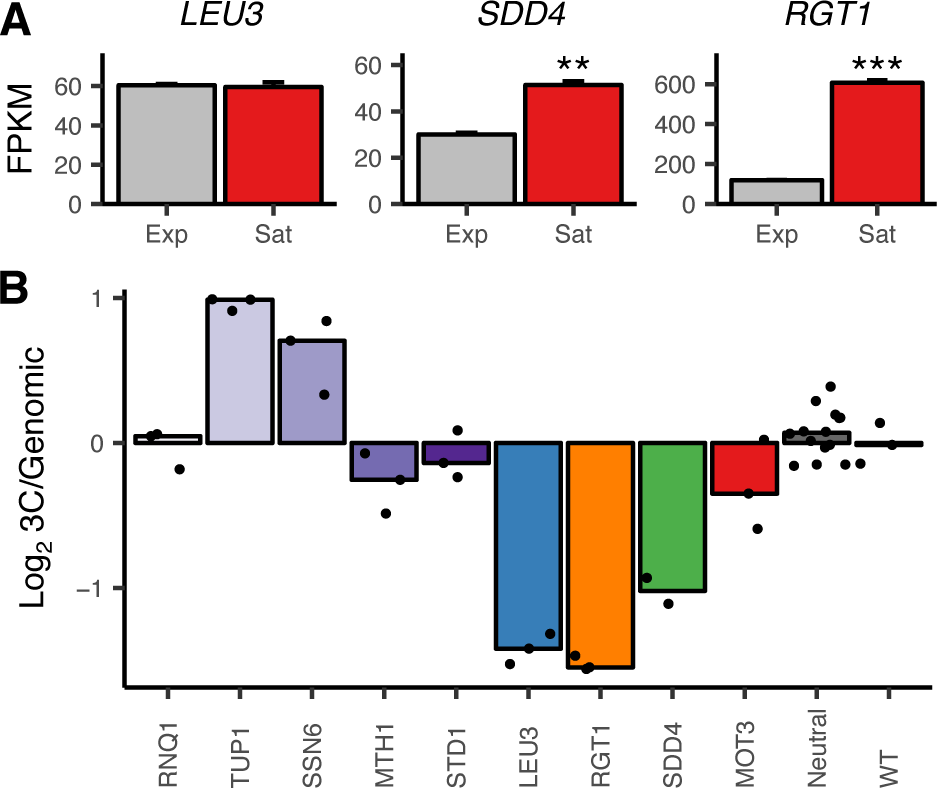
**Transcription factor expression levels and the Tup1/Ssn6 repressor complex regulate *HAS1pr-TDA1pr* pairing.** (A) RNA-seq expression levels for *LEU3*, *SDD4*, *RGT1* in exponentially growing (Exp) and saturated (Sat) cultures. Bars indicate mean ± s.e.m. of biological triplicates. \*\**P* < 0.01, \*\*\**P* 0.001 (Student’s t-test). (B) Bar plot of median relative abundance in 3C library compared to the genomic library, with overlaid scatter plot of individual barcoded deletion strains. Retested controls are shown in same colors as in **Figure 2**; potential interaction partners are shown in shades of purple.

Rgt1 is known to interact with several cofactors that affect its DNA binding and transcriptional repression activities: the Tup1/Ssn6 co-repressor complex and the proteins Mth1 and Std1 (Polish et al., 2005). Our experiments thus far had not distinguished whether Rgt1’s pairing activity is directly mediated by physical interactions among molecules of Rgt1 or indirectly, through these or other interaction partners. To address this, we performed *trans* MAP-C for these four interacting partners of Rgt1, along with the same positive and negative controls as before (**Figure 3B**). In addition, based on the glutamine and asparagine-rich domains present in Sdd4 and Rgt1, we tested deletion of *RNQ1*, a Q/N-rich peptide known to influence the oligomerization of other Q/N-rich proteins (Derkatch et al., 2004). Deletion of *RNQ1* had no effect on pairing, suggesting that *HAS1pr-TDA1pr* pairing is not mediated by Q/N-rich domains. Deletion of *TUP1* or *SSN6* both led to increased pairing, indicating that the recruitment of the Tup1/Ssn6 co-repressor complex inhibits pairing. This is consistent with its known inhibition of Rgt1’s DNA binding activity (Roy et al., 2013), which is required for pairing. Deletion of *MTH1* or *STD1* had a minimal negative effect on pairing. Taken together, these data suggest that the role of Rgt1 in pairing is not simply to recruit cofactors that mediate pairing; instead, its pairing activity may compete with its transcriptional repression activity.

### A combination of transcription factors mediates pairing

We next explored whether a combination of transcription factors is necessary for pairing, using overexpression and analyses of genome-wide chromosome conformation data. Based on our inability to find any protein other than the transcription factors Leu3, Sdd4, and Rgt1 required for pairing in saturated culture conditions, we wondered whether overexpression of any one of the three proteins would be sufficient to produce pairing in non-saturated culture conditions. We used the Z_3_EV estradiol induction system (McIsaac et al., 2014) to individually overexpress Leu3, Sdd4, or Rgt1, and measured pairing between the native *HAS1pr-TDA1pr* loci using 3C in *S. cerevisiae* x *S. uvarum* hybrids (**Figure 4A**). In all three strains, a 2 h estradiol induction led to no increase in pairing strength. As an alternative test, we used galactose induction to overexpress *RGT1*, and observed a decrease in pairing strength relative to a strain lacking the overexpression cassette (**Figure S6**). These results are consistent with no single TF being sufficient for pairing; however, Leu3 and Rgt1 are both known to change in conformation (Sze et al., 1992) or phosphorylation state (Kim et al., 2003) in different conditions, so it remains possible that overexpression of a single TF in the correct state suffices for pairing.

**Figure 4.**
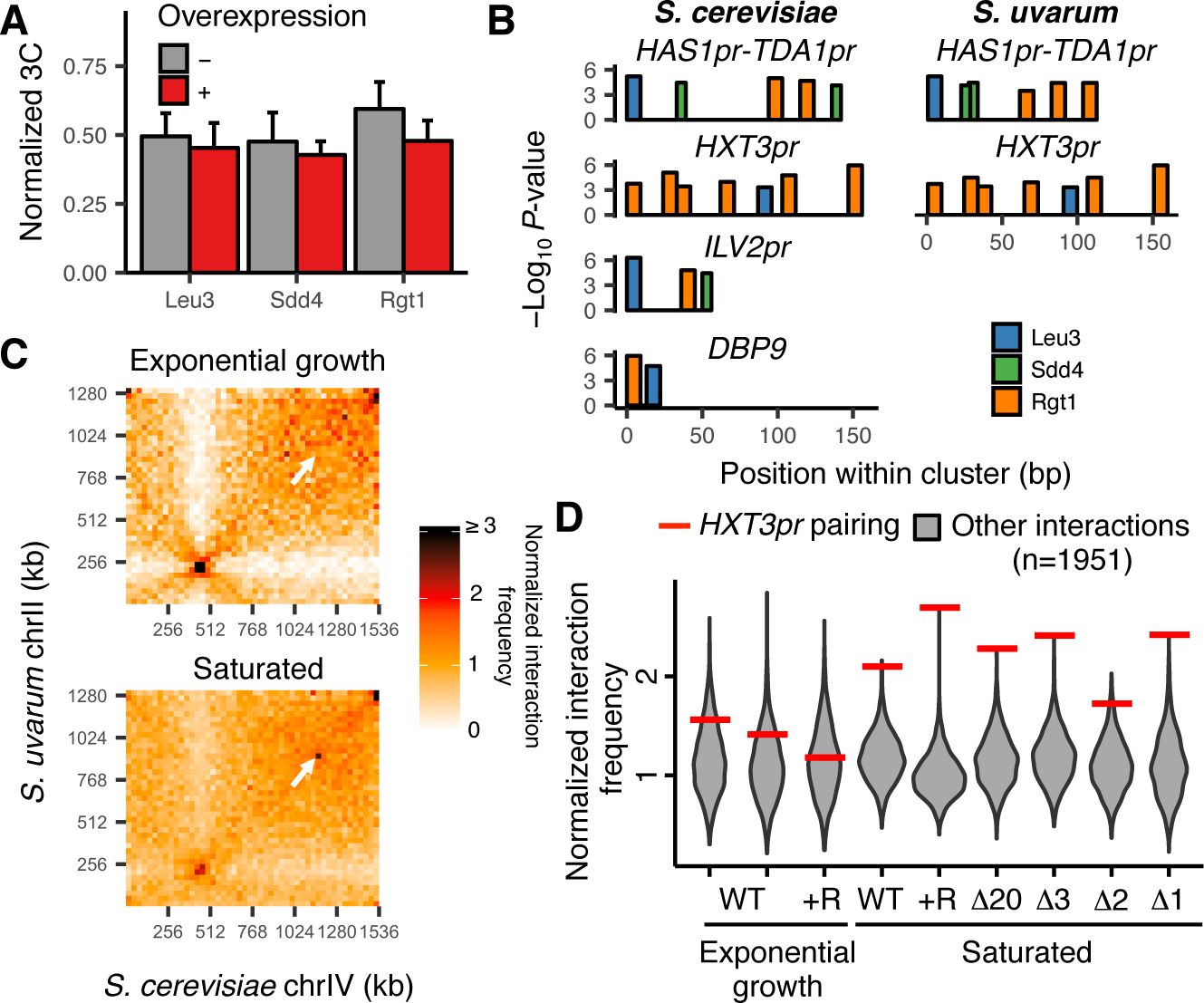
**Combinatorial TF binding mediates inducible interchromosomal contacts.** (A) 3C for *HAS1pr-TDA1pr* homolog pairing with and without estradiol-induced overexpression of *LEU3*, *SDD4*, *RGT1* in *S. cerevisiae* x *S. uvarum* hybrids, normalized to contacts between *HAS1pr-TDA1pr* and *LCB1* on *S. cerevisiae* chrXIII. Bars indicate mean ± s.d. of technical triplicates. (B) Top clusters of Leu3, Sdd4, and Rgt1 motifs containing Leu3 and Rgt1 motifs in *S. cerevisiae* and *S. uvarum* genomes, in order of lowest *P*-value from top to bottom. (C) Hi-C contact maps of *S. cerevisiae* chrIV interactions with *S. uvarum* chrII at 32 kb resolution in exponentially growing and saturated cultures of a *S. cerevisiae* x *S. uvarum* hybrid strain (YMD3920). White arrows indicate interactions between homologous *HXT3* promoters. (D) Strength of *HXT3* promoter pairing across conditions and strain backgrounds, at 32 kb resolution (red lines) compared to similar interactions (grey violin plots; i.e. interactions between an *S. cerevisiae* locus and an *S. uvarum* locus, where both loci are ≥15 bins from a centromere and ≥2 bins from a telomere, ≥2 bins from *HAS1pr-TDA1pr*, and not both on chrXII). WT represents strain ILY456. +R indicates a restriction site added upstream of *HAS1* (YMD3920), ∆20 indicates a 20 kb deletion centered at *S. cerevisiae HAS1* (YMD3266), ∆3 indicates a 3 kb deletion centered at *S. cerevisiae HAS1* (YMD3267), ∆2 indicates a 2 kb deletion of the *S. cerevisiae HAS1* coding sequence (YMD3268), and ∆1 indicates a 1 kb deletion of the *S. cerevisiae HAS1pr-TDA1pr* intergenic region (YMD3269).

If the clustered binding of Leu3, Sdd4, and Rgt1 is necessary and sufficient to cause pairing in saturated culture conditions, either 1) no loci other than *HAS1pr-TDA1pr* have all three TFs bound nearby, or 2) other loci that do have all three TFs bound nearby should also exhibit pairing. To test the first possibility, we scanned the *S. cerevisiae* and *S. uvarum* genomes for clusters of the three motifs. Indeed, *HAS1pr-TDA1pr* was the only locus with matches for all three motifs in both the *S. cerevisiae* and *S. uvarum* genomes (**Figure 4B**). A second locus upstream of *ILV2* also had all three matches, but only in the *S. cerevisiae* genome. This suggests that the strong homolog pairing at the *HAS1pr-TDA1pr* is explained by its unique juxtaposition of transcription factor binding sites.

Our *trans* knockout screen suggested that *SDD4* plays a weaker role in pairing than *LEU3* or *RGT1*; therefore, we expanded our search to motif clusters with only Leu3 and Rgt1 motifs. This revealed a second locus, upstream of *HXT3*, which contains both Leu3 and Rgt1 motifs in both the *S. cerevisiae* and *S. uvarum* genomes (one other locus, in *DBP9*, contained both motifs only in the *S. cerevisiae* genome) (**Figure 4B**). We wondered whether the *HXT3* promoters pair inducibly like the *HAS1pr-TDA1pr* locus. To address this question, we leveraged our previously published Hi-C datasets (Kim et al., 2017), along with new Hi-C data for the background in which we performed our pooled mutant experiments, to compare the strength of *HXT3pr* pairing to other interchromosomal interactions and assess whether this pairing is condition-specific. Indeed, across several strain backgrounds, the *HXT3* promoters exhibited 1.6- to 2.7-fold increased interaction frequencies compared to other similar interchromosomal pairs of loci (at least 480 kb from a centromere and excluding subtelomeric regions) in saturated culture conditions (**Figure 4C and D**), albeit weaker than the pairing between *HAS1pr-TDA1pr* (**Figure S7**). This pairing occurred only at baseline levels during exponential growth in rich medium. These results suggest that Leu3 and Rgt1 act together to mediate inducible interchromosomal homolog pairing not only at the *HAS1pr-TDA1pr* locus, but also at the *HXT3* promoter.

The identification of two pairs of loci, *HAS1pr-TDA1pr* and *HXT3pr*, exhibiting homolog pairing mediated by Leu3 and Rgt1 opened the possibility that there might be cross-pairing between *HAS1pr-TDA1pr* and *HXT3pr*. To test this possibility, we extracted the interaction frequencies among the four Leu3- and Rgt1-bound loci from our Hi-C data. In all saturated culture datasets where homolog pairing was present, inter-locus interactions were substantially weaker than homolog pairing interactions and similar in frequency to those in non-pairing conditions (**Figure S7**). Together with our previous experiment demonstrating that two identical copies of the minimal pairing region can pair even at non-allelic locations in haploid *S. cerevisiae* (**Figure 1C**), these data suggest that the interchromosomal pairing mediated by Leu3 and Rgt1 is sequence-specific beyond the simple presence of the same TF binding sites.

## Discussion

In summary, we developed Mutation Analysis in Pools by Chromosome conformation capture (MAP-C), a method to simultaneously test hundreds of mutations for their effects on a chromosomal contact of interest. MAP-C can be used to identify the precise sequences that are necessary for a contact (*cis* MAP-C) as well as the factors that are necessary to mediate the contact (*trans* MAP-C). Here we applied both flavors of MAP-C to dissect the mechanism of inducible interchromosomal pairing between *HAS1pr-TDA1pr* alleles in budding yeast. Using a combination of gain-of-function and loss-of-function screens, we demonstrate that a trio of transcription factors—Leu3, Sdd4, and Rgt1—mediate pairing between clusters of binding sites. Clusters of all three TF motifs are rare, and are conserved between the diverged *S. cerevisiae* and *S. uvarum* genomes at only the *HAS1pr-TDA1pr* locus; however, the *HXT3* promoter, which contains Leu3 and Rgt1 motifs, also exhibits inducible pairing in saturated cultures.

Our results begin to elucidate the mechanisms of condition-specific interchromosomal contacts and homolog pairing, which have been elusive (Mirkin et al., 2013). Unlike more prevalent nuclear-pore mediated gene relocalization and homolog pairing (Brickner et al., 2012; Randise-Hinchliff and Brickner, 2016), *HAS1pr-TDA1pr* pairing in saturated cultures appears not to require the nuclear pore complex (**Figure 2B**). We have not yet fully defined the biochemical mechanism of pairing—it is possible the TFs interact directly and/or indirectly—but so far, no known interaction partners have proven essential for pairing. Instead, we have found that the Tup1/Ssn6 repressor complex recruited by Rgt1 negatively regulates pairing, perhaps by inhibiting the DNA-binding activity of Rgt1. This suggests the repressor activity of Rgt1 competes with its pairing activity; more studies will be needed to carefully map the relationship between the different functions of Rgt1.

Another question is whether the contacts are stoichiometric, *e.g.* one-to-one, or instead mediated by aggregation of one or more TFs via weak interactions. Notably, we observe homotypic pairing between homologs of *HAS1pr-TDA1pr* and *HXT3pr* but not heterotypic pairing, suggesting that Leu3 and Rgt1 do not indiscriminately form contacts between all of their binding sites. Instead, only pairs of loci with a series of motifs in similar orders and orientations form frequent contacts, consistent with a stoichiometric interaction model. This added specificity beyond the presence of TF motifs could explain the lack of pairing between *HAS1pr-TDA1pr* and other clusters of Leu3 and Rgt1 motifs, *e.g. ILV2pr* and *DBP9*. We hypothesize that the globally acting chromosome conformation effects of the cohesin and condensin complexes (Lazar-Stefanita et al., 2017; Schalbetter et al., 2017) are complemented by highly condition-specific and localized interactions coordinated by combinations of TFs.

Although our studies were concentrated on *HAS1pr-TDA1pr* pairing in budding yeast, our method should be applicable to other loci and organisms. The main constraints in experimental design are that 1) the introduced mutations or associated barcodes must be included in the 3C PCR product, and 2) the region of interest should not be digested by the restriction enzyme. As we have implemented this approach, the region targeted by saturation mutagenesis was limited to 250 bp to allow for Illumina sequencing, but this could be extended using either barcode association, similar to our *trans* knockout screens, or long-read sequencing methods. Also, we focused on a single pair-wise interaction, but it is also possible to assay a mutant pool for multiple interactions, by using multiple primer pairs. MAP-C should be applicable to intrachromosomal contacts as well as interchromosomal ones, albeit with a potentially higher background of nonspecific contacts.

MAP-C leverages the high throughput of saturation mutagenesis and mutant collections to allow systematic dissection of chromosome conformation. We have tested up to ~1,000 variants at a time, but with larger-scale experiments, it should be possible to test even more variants. A major potential strength of our approach is that unlike cellular high-throughput genetic screens (Fowler and Fields, 2014; Gasperini et al., 2016; Shalem et al., 2015), it resolves the functional consequences of mutations at the allelic level, and thus is not confounded by heterozygosity (Patwardhan et al., 2009). As we continue to map chromosome conformation at high resolution across ever-expanding numbers of cell types and conditions, MAP-C will provide a scalable approach to dissect the molecular mechanisms underlying specific contacts.

## Acknowledgments

We thank Katherine Xue, William Noble, and members of the Shendure and Dunham labs for comments and discussion, Ivan Liachko for help with Hi-C experiments, Noah Hanson for help with yeast deletion collections, and Yixian Zheng, Monica Sanchez, and R. Scott McIsaac for strains. This work was supported by grants from the NIH (U54 DK107979 to J.S.) and NSF (graduate research fellowship DGE-1256082 to S.K.). M.J.D. is a Senior Fellow in the Genetic Networks program at the Canadian Institute for Advanced Research and is supported in part by a Faculty Scholar grant from the Howard Hughes Medical Institute. J.S. is an investigator of the Howard Hughes Medical Institute.

## Author Contributions

Conceptualization, S.K.; Methodology, S.K.; Investigation, S.K.; Writing - Original Draft, S.K.; Writing - Review & Editing, S.K., M.J.D., and J.S.; Visualization, S.K.; Supervision, M.J.D. and J.S.

## Materials and Methods

### Yeast Strains and Culture

Yeast strains used in this study are listed in Table S1. Yeast were cultured at 30C, with the exception of *S. uvarum* strains, which were grown at room temperature. Cultures were grown shaking overnight to OD_600_ > 5 for saturated culture samples, or diluted to OD_600_ ~ 0.125 and grown to OD_600_ = 0.5-0.8 for exponential growth samples. Estradiol inductions were performed by addition of beta-estradiol to 1 uM final concentration (or equivalent volume of ethanol for negative control) to OD_600_ = 0.5 cultures grown in YPD (1% w/v yeast extract, 2% w/v peptone, 2% w/v dextrose) and grown for 2 hours. Galactose induction was performed by growth in synthetic complete medium (without uracil for selection for overexpression plasmid) with 2% v/v raffinose to OD_600_ = 0.75 followed by addition of 2% galactose and subsequent growth for 1.5 hours. For comparison, yeast were grown in synthetic complete medium with or without uracil with 2% glucose. *S. cerevisiae* x *S. uvarum* hybrids were generated by standard mating and auxotrophic or drug selection procedures. Yeast transformations were performed using a modified Gietz LiAc method (Pan et al., 2004).

### Mutant Library Generation

*Subsequences.* All 178 bp subsequences of the intergenic region between the *S. cerevisiae HAS1* and *TDA1* coding sequences were synthesized in an array-synthesized oligonucleotide pool, and then amplified and cloned into a vector just downstream of a KanMX cassette followed by a DpnII restriction site (GATC) using NEBuilder HiFi. These plasmids, which contain homology to the *S. cerevisiae HAS1* and *TDA1* sequences, were linearized by restriction digestion and transformed into YMD3919 (deletion of *S. cerevisiae has1pr-tda1pr*). Transformants (typically ~5,000 per experiment) were selected on G418 medium for a total of at least 4 days (including two rounds of scraping and replating a portion onto a fresh plate), and then used to inoculate a 50 ml culture in YPD (1% w/v yeast extract, 2% w/v peptone, 2% w/v dextrose).

*Error-prone PCR.* The primers TTCACCGCCTGCTATCATCC and GAATCGGCGGAATAACCTAACACG were used in a PCR reaction using the Agilent GeneMorph II kit, using as template 0.1 ng of a PCR product generated with the same primers. The error-prone PCR products were then cloned, transformed, and selected as described above.

*Programmed 3 bp substitutions.* For each set of 3 consecutive base-pairs between positions 532-675 (inclusive), we randomly chose three trinucleotide substitutions such that among them, each nucleotide change (*e.g.* A->C) was included once at each position and so that no DpnII restriction sites (GATC) were created. These sequences were synthesized on an array and processed as described above.

*In-gene knockout screen.* The selected gene knockout strains from the MATalpha yeast deletion collection, in addition to the knockouts of the three nearest genes on either side excluding those absent or known to be slow-growing in the yeast deletion collection, were pooled and transformed *en masse* using a PCR construct with homology arms for the TEFb promoter and terminator from the KanMX deletion cassette flanking the pairing sequence (**Figure S1B**), an EcoRI restriction site, and the *URA3* selectable marker gene. The entire pool was selected on synthetic complete medium without uracil and processed as described above.

*Fixed-locus knockout screens.* TF genes were defined as genes with motifs on YeTFaSCo (de Boer and Hughes, 2012) that are not annotated as being part of a complex. Genes with no non-systematic name in the GFF file from the Saccharomyces Genome Database (SGD; version R64.2.1) were excluded. In addition, we included knockouts of genes described as nuclear pore components in SGD, the wild type strain and knockouts of eight genes known to have minimal fitness consequences (Payen et al., 2016). The selected gene knockout strains from the MATa yeast deletion collection (excluding those failing quality control or known to grow slowly) were grown in separate wells of deep 96-well plates, and transformed in 96-well format (**SI Materials and Methods**) with a PCR construct similar to that used in the in-gene knockout screen, but with a unique 12 bp barcode added upstream of the pairing sequence, and homology to the *YDR535C* coding sequence. Colonies were picked and verified by PCR, and one successful clone from each strain was pooled together and then diluted and grown overnight in YPD. Colonies from positive and negative control strains were repooled with new strains for the validation experiment.

*Interactor knockout screen*. The selected gene knockout strains from the MATa yeast deletion collection were each transformed with a cocktail of constructs with 14 different barcodes each, and then at least 8 colonies were Sanger sequenced, and three colonies from each strain carrying different barcodes were pooled and processed as described above.

### 3C

Cells were crosslinked by addition of 37% formaldehyde to a final concentration of 1% (v/v) and incubation at room temperature for 20 min, quenched by addition of 2.5M glycine to a final concentration of 150 mM and incubation at room temperature for 5 min, and then washed in 1x Tris-buffered saline (TBS) and stored as a pellet at −80C in aliquots of 50-100 ul dry pellets. Cells were lysed by vortexing in lysis buffer (TBS + 1% Triton X-100 with protease inhibitor cocktail (Pierce)) with 500 um glass beads for 6 cycles of 2 min, with 2 min on ice between cycles. The lysate was collected by puncturing the bottom of each tube and then centrifuging the tube, stacked on top of an empty tube. The lysate was then washed in lysis buffer, then TBS, and finally resuspended in 10 mM Tris pH 8.0 to a volume of ~200 ul per 25 ul of starting dry pellet volume. A single 200 ul aliquot of lysate was then precleared by addition of 0.2% SDS and incubation at 65C for 10 min, cooled on ice, quenched by addition of 1% Triton-X100 (v/v), and then digested overnight with at least 100U of restriction enzyme (200U DpnII for *cis* experiments and galactose induction, 400U EcoRI-HF for *trans* experiments, and 100U NlaIII for estradiol inductions). The restriction digest was heat-inactivated at 65C for 20 min in the presence of 1.3% SDS, and then chilled on ice and added to a dilute ligation reaction in 4 ml volume with 1% Triton-X100, 1x T4 DNA Ligase Buffer (NEB), and 10,000U of T4 DNA ligase (NEB) and incubated at room temperature for 4 hours. The ligation products were reverse-crosslinked with proteinase K at 65C overnight, and then purified by phenol-chloroform extraction followed by clean-up on a Zymo DNA Clean & Concentrator-5 column. The resulting 3C DNA was quantified using a Qubit. Each technical replicate was processed separately beginning with cell lysis.

### Library Preparation and Sequencing

3C libraries were prepared by amplification of up to 8 reactions of 50 ng 3C DNA per replicate, using primer pairs specific to the chromosomal contact of interest (for pairing library) or a control off-target chromosomal contact (for off-target library), for 24-32 cycles. Genomic libraries were prepared by amplification of up to 4 reactions of 50 ng of either 3C DNA or genomic DNA using primers flanking the targeted mutations or barcodes, for 17-22 cycles. Reactions for each replicate were pooled, purified by Ampure XP beads, and then re-amplified with primers flanking the mutagenized or barcode region and including sequencing adapter sequences for 5-8 cycles, and then again with primers adding sample indices and Illumina flow-cell adapters for 6-9 cycles. All reactions were prepared with KAPA HiFi HotStart ReadyMix with recommended thermocycling conditions, and included 0.5x SYBR Green I to monitor amplification by quantitative PCR and minimize the number of PCR cycles. The final libraries were sequenced on an Illumina MiSeq or Nextseq 500 using paired-end sequencing. See Table S2 for detailed information on each library.

### Sequencing Analysis

Paired-end reads were merged and adapter-trimmed using PEAR (Zhang et al., 2014), except for *trans* knockout experiments, in which only read 1 was used. These reads were then trimmed of the first 4 bp (corresponding to a randomized region for Illumina clustering purposes) and mapped using Bowtie 2 (Langmead and Salzberg, 2012).

*Subsequences*. Reads were mapped to the *S. cerevisiae HAS1pr-TDA1pr* region, and then the read coverage was calculated using bedtools (Quinlan and Hall, 2010).

*Error-prone PCR.* Reads were mapped to the wild-type sequence of the mutagenized region. The resulting alignments were scored for number of substitutions, insertions, and deletions, and the fraction of reads with a substitution at each position were calculated.

*3 bp substitutions*. Reads were mapped to the wild-type sequence of the mutagenized region. The resulting alignments were scored for number of substitutions, insertions, and deletions, and the fraction of reads with a substitution at each position were calculated, excluding reads with fewer than 3 substitutions (which correspond to PCR or sequencing errors of the wild-type sequence).

*Trans knockout screens*. Reads were mapped to all 192 possible barcode sequences, and the normalized fraction of reads mapping to a given barcode with a MAPQ ≥ 20 in the 3C library compared to the genomic control was calculated for each replicate.

### 3C qPCR

3C DNA was amplified using the same conditions as MAP-C libraries, in three replicates per primer pair. The pairing 3C products were normalized to the off-target (intrachromosomal) 3C products, assuming 2-fold amplification per cycle.

### Motif Analysis

Motif clusters were identified using MCAST (Grant et al., 2015) with the default motif p-value threshold of 0.0005, and an E-value threshold of 2, using the high-confidence motifs for Leu3 (#781), Rgt1 (#2227), and Sdd4 (#588) from YeTFaSCo (de Boer and Hughes, 2012). Individual motif occurrences for each TF were scored for the *S. cerevisiae* (version R64.2.1) and *S. uvarum* genomes (as revised in (Kim et al., 2017)) using FIMO (Grant et al., 2011) with the default p-value threshold of 0.0005 and the option --max-strand.

### Hi-C

Hi-C was performed and analyzed as in (Kim et al., 2017) using the restriction enzyme Sau3AI.

### Data availability

All sequencing data have been deposited in the Gene Expression Omnibus (GEO) under accession number GSE118118. Hi-C and RNA-seq data from **Figure 4** are from GEO accession number GSE88952.

## Supplementary Information

### SI Materials and Methods

#### 96-well yeast transformation

##### Day 1

1. Array strains into shallow 96-well plate, resuspending one colony of each strain in 100 ul YPD.

2. Incubate overnight at 30C.

##### Day 2

3. Transfer 45 ul into each of two deep 96-well plate with 400 ul YPD per well. One plate will be used to make glycerol stocks.

4. Incubate overnight at 30C.

##### Day 3

5. With one deep 96-well plate, make glycerol stocks by resuspending each well by pipetting and then transferring 80 ul to a U-bottom 96-well plate with 40 ul of 50% glycerol already added. Store at −80C.

6. With the other deep 96-well plate, carefully remove 360 ul (180 ul twice) supernatant by pipetting.

7. Add 200 ul 2xYPAD and resuspend by pipetting.

8. Incubate at 30C for 1.5 hrs.

9. During this incubation, denature salmon sperm DNA (ssDNA) by placing in 98C heat block for 5 min, and then place on ice.

10. Pellet by centrifuging at 2000g for 2 min.

11. Remove 200 ul supernatant by pipetting.

12. Add 180 ul water and resuspend by pipetting.

13. Pellet by centrifuging at 2000g for 2 min.

14. Remove supernatant by pipetting.

15. Add 180 ul 0.1M LiAc and resuspend by pipetting.

16. Pellet by centrifuging at 2000g for 2 min.

17. Remove supernatant by pipetting

18. Add 120 ul 50% PEG-3350 + 18 ul 1.0M LiAc + 5 ul denatured ssDNA (pre-mixed together: 26 ml 50% PEG-3350, 3.9 ml 1.0M LiAc, 1.083 ml ssDNA; use 143 ul per well)

19. Resuspend by pipetting.

20. Add 20 ul DNA, mix by pipetting up and down.

21. Resuspend by gently vortexing.

22. Incubate at 30C for 30 min.

23. Resuspend by gently vortexing.

24. Incubate at 42C for 40 min in water bath.

25. Pellet by centrifuging at 2000g for 2 min.

26. Remove 180 ul supernatant by pipetting.

27. Add 160 ul water, resuspend by pipetting

28. Pellet by centrifuging at 2000g for 2 min

29. Remove supernatant by pipetting

30. Add 100 ul water, and resuspend by pipetting.

31. Add 1 ml C-ura+2% glucose. Pipette up and down to mix.

32. Incubate at 30C overnight.

##### Day 4

33. Remove supernatant by pipetting.

34. Add 50 ul water, resuspend by pipetting/shaking, transfer to shallow 96-well plate

35. Pin/pipette onto C-ura+2% glucose plate(s)

36. Incubate at 30C for 2-4 days

37. Pick colonies for PCR verification and grow in shallow 96-well plate with 150 ul C-ura+2% glucose

## Supplementary Tables

**Table S1.**
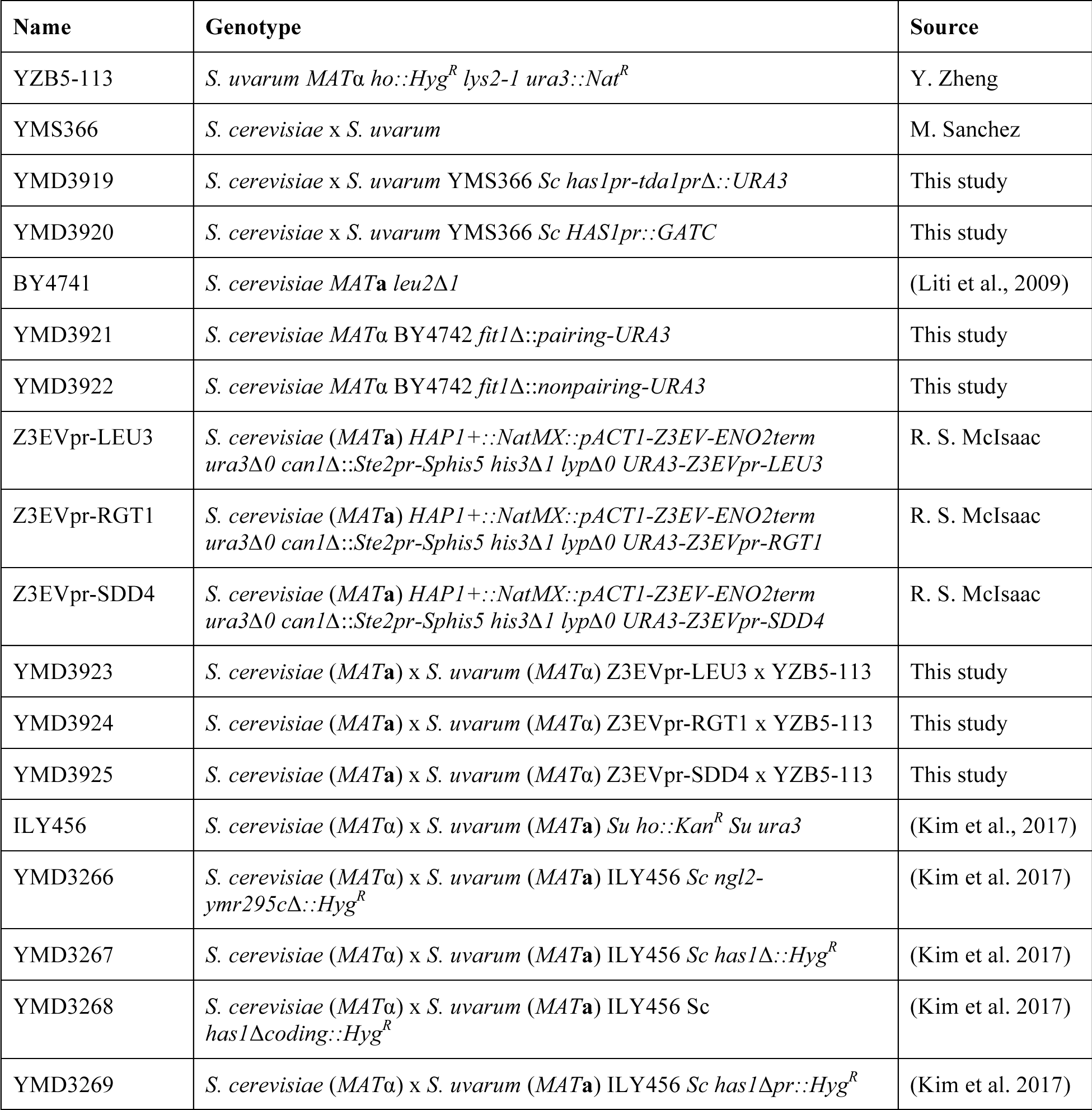
Yeast strains used in this study.

**Table S2. Table of experiments and conditions**

## Supplementary Figures

**Figure S1.**
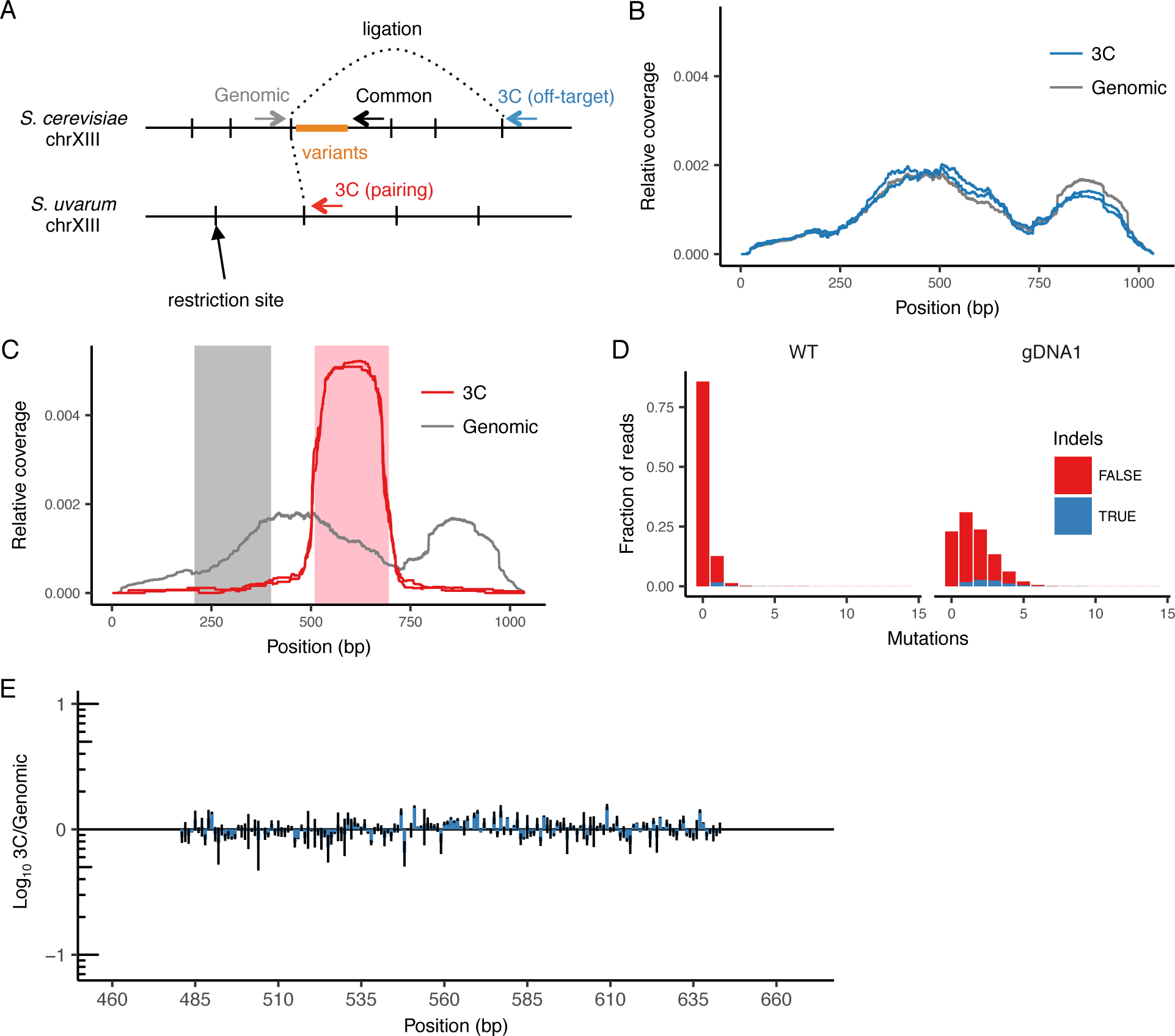
**Design and controls for using *cis* MAP-C to dissect *HAS1pr-TDA1pr* pairing.** (A) Schematic (not to scale) of primer locations for *cis* MAP-C experiments. (B) Off-target control results for 178 bp subsequences of *S. cerevisiae HAS1pr-TDA1pr*, shown as read coverage for 3C and genomic libraries. Each line represents a technical replicate. (C) Regions of *HAS1pr-TDA1pr* used for testing ectopic pairing. Pairing sequence shown in pink, and non-pairing control shown in gray. (D) Distribution of the number of mutations in the wild type and error-prone PCR genomic libraries. (E) Off-target control results for error-prone PCR mutagenesis of central *S. cerevisiae HAS1pr-TDA1pr*, shown as ratio of the total substitution frequency at each position in the 3C library compared to the genomic library. Error bars indicate the two technical replicates.

**Figure S2.**
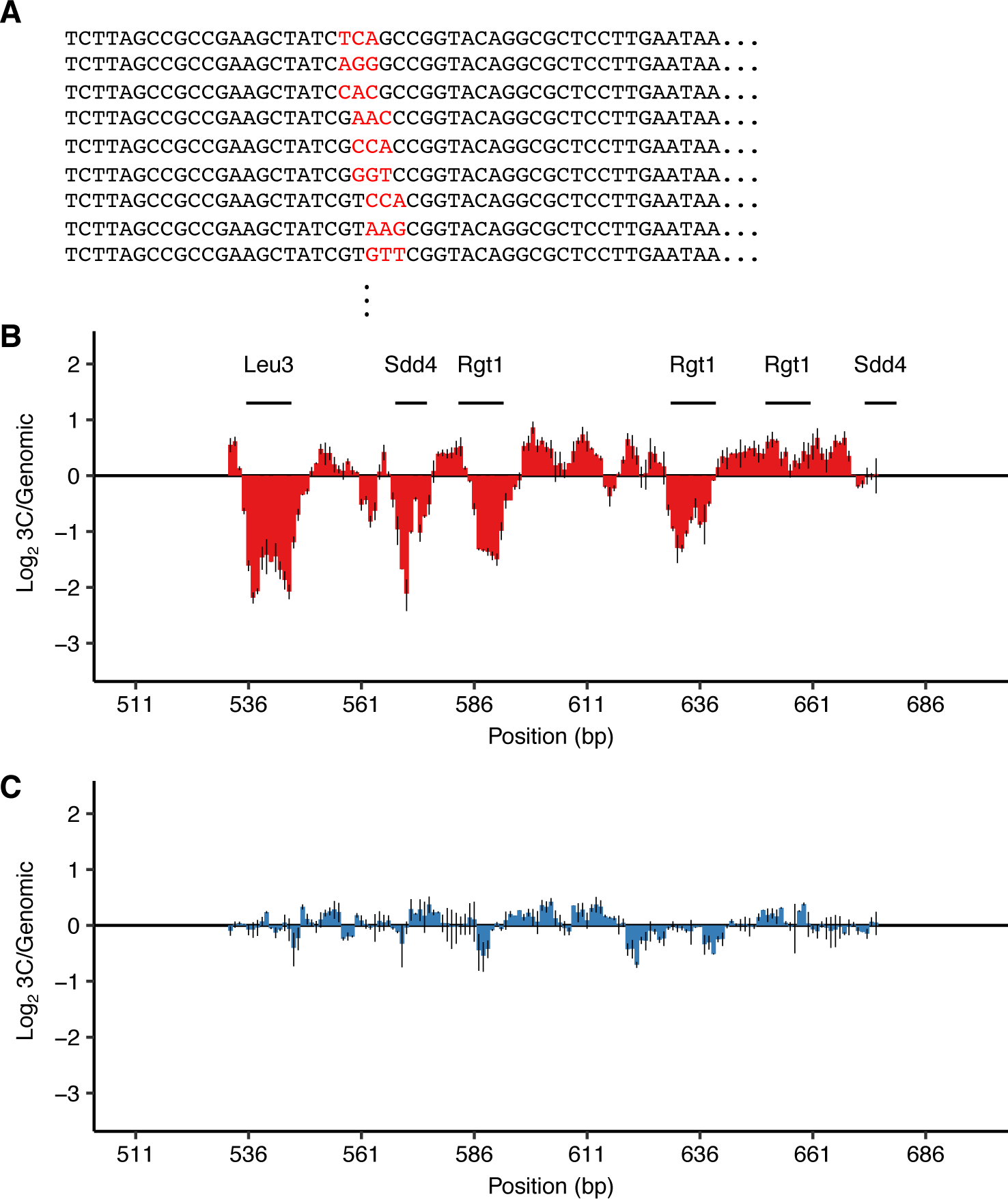
**Validation of TF motifs required for *HAS1pr-TDA1pr* pairing with a 3 bp substitution mutant library.** (A) Library design. Examples of mutant sequences, with mutations in red. Each set of 3 bp was replaced with three random 3 bp sequences so that every nucleotide was switched to every other nucleotide. (B-C) Ratio of the total substitution frequency at each position in the 3C library compared to the genomic library, for contacts between *S. cerevisiae HAS1pr-TDA1pr* and the *S. uvarum* homolog (B) or *S. cerevisiae LCB1*, an off-target control (C). Horizontal bars indicate the positions of TF motif matches.

**Figure S3.**
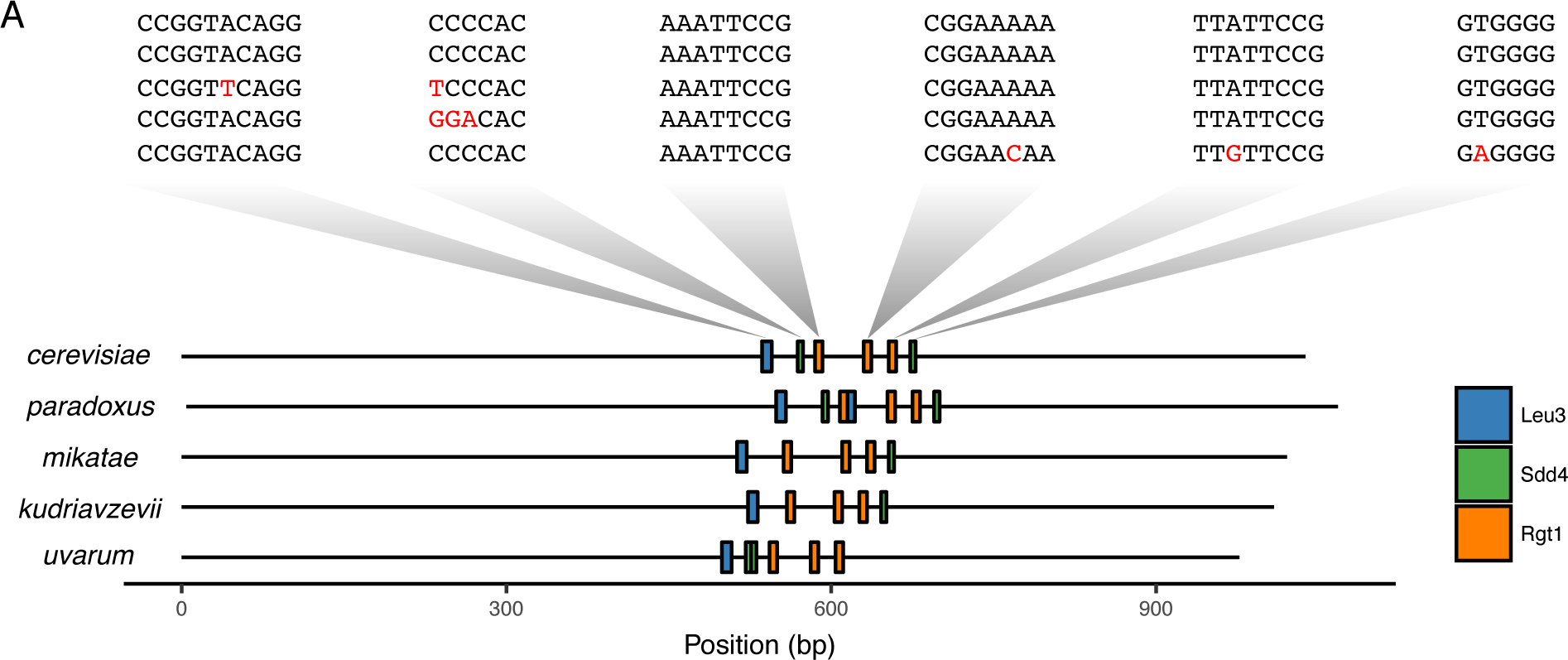
**Conservation of TF motifs in *HAS1pr-TDA1pr*.** Position of Leu3 (CCGNNNN(A/C)GG), Rgt1 (CGGAA), and Sdd4 (CCCCAC) motifs in the intergenic region between *HAS1* and *TDA1* in *Saccharomyces sensu stricto* yeasts *S. cerevisiae*, *S. paradoxus*, *S. mikatae*, *S. kudriavzevii*, and *S. uvarum* (order of divergence from *S. cerevisiae*). Above, the sequences of the regions aligning to the *S. cerevisiae* motifs are shown in the same vertical order, with mismatches in red.

**Figure S4.**
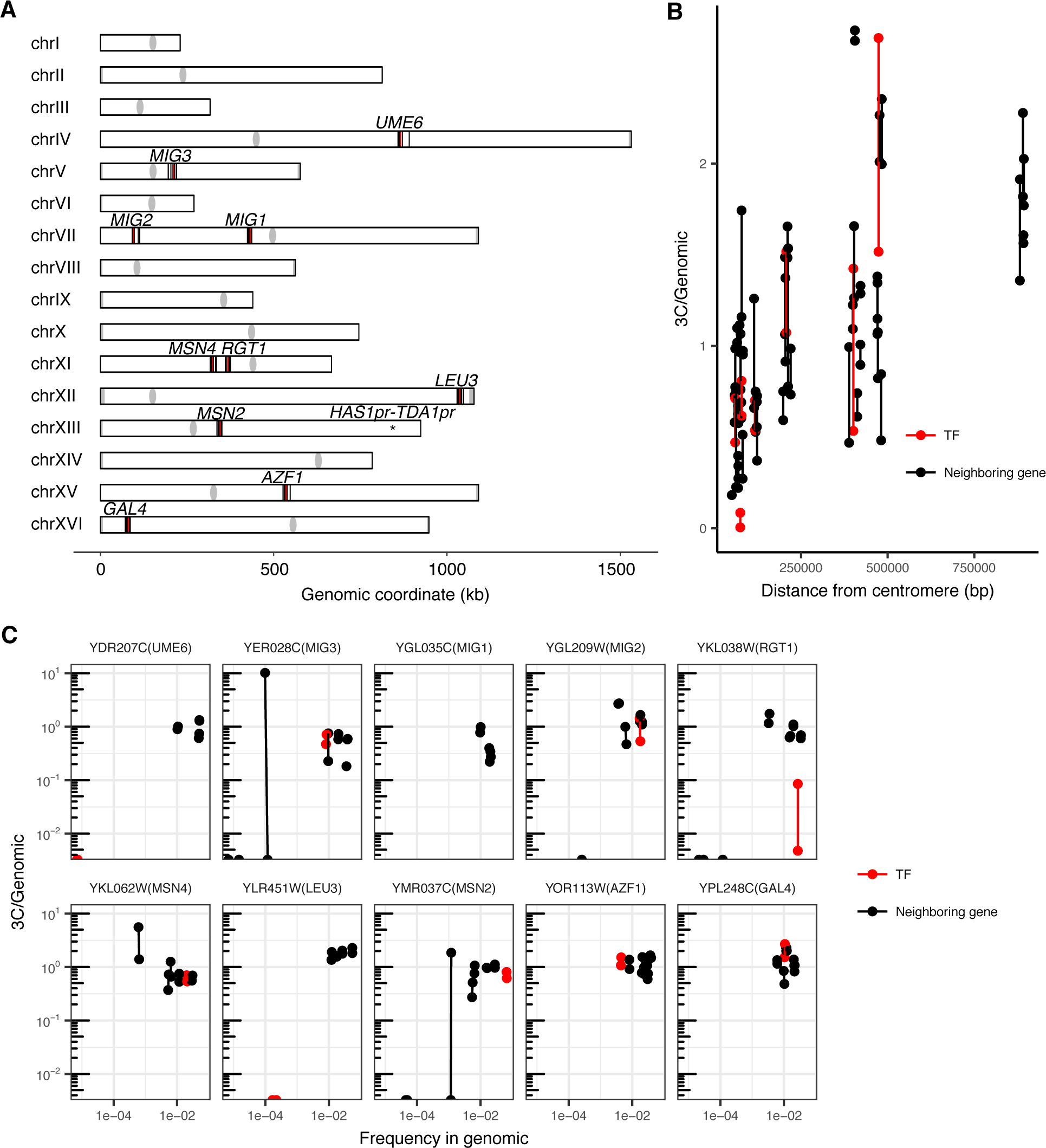
**A pilot TF gene knockout screen for *HAS1pr-TDA1pr* pairing.** (A) Genomic locations of the tested TF gene knockouts (in red) and tested genomic neighbors (in black). (B) Effect of centromeric distance on ectopic *HAS1pr-TDA1pr* pairing in haploid *S. cerevisiae*. Each pair of connected dots represent two technical replicates. (C) Ratio of barcoded strain abundance in 3C library vs. genomic library for each of 10 tested TF gene knockouts, along with genomic neighbors.

**Figure S5.**
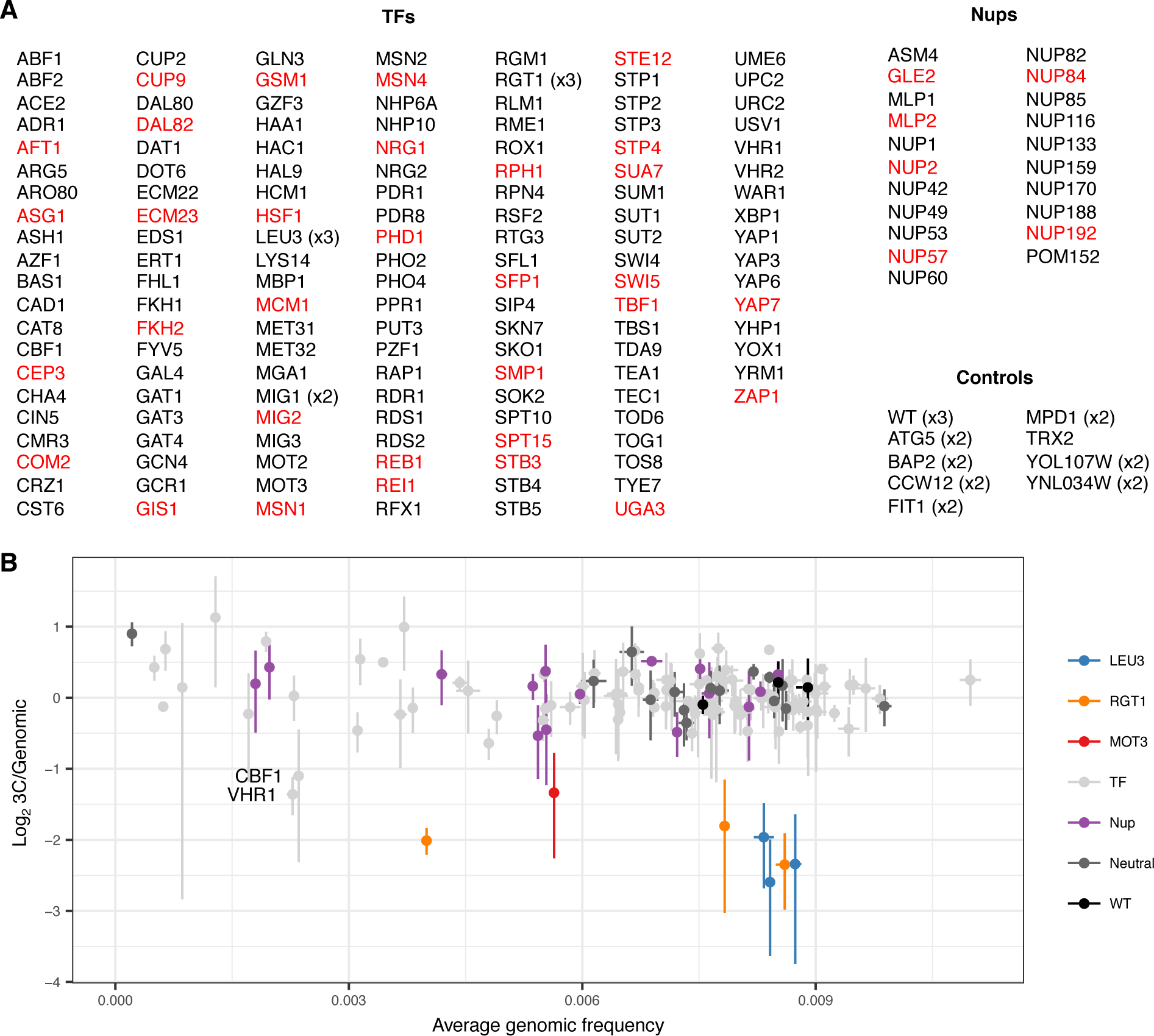
**An expanded *trans* knockout screen for *HAS1pr-TDA1pr* pairing.** (A) List of tested gene knockouts. Strains that dropped out of the pool during library construction are shown in red. (B) Scatter plot of each barcoded strain’s abundance in the genomic library vs. ratio of abundance in the 3C library compared to the genomic library. Center values indicate mean; error bars indicate s.d. of three technical replicates.

**Figure S6.**
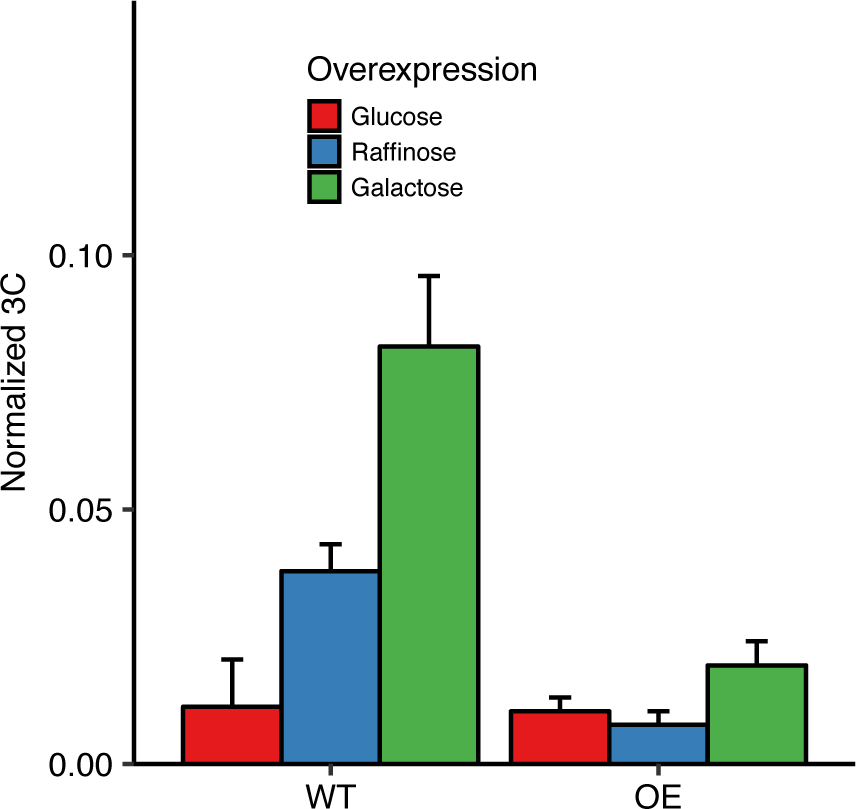
**Overexpression of *RGT1* is not sufficient for *HAS1pr-TDA1pr* pairing.** 3C for contact between *HAS1pr-TDA1pr* alleles normalized to the frequency of contacts between *S. cerevisiae HAS1pr-TDA1pr* and *S. cerevisiae LCB1* in *S. cerevisiae* x *S. uvarum* hybrids with (OE) or without (WT) a plasmid carrying a copy of *RGT1* under a *GAL* promoter, grown in glucose (red), raffinose (blue), or galactose (green). Bars indicate mean ± s.d. of technical triplicates.

**Figure S7.**
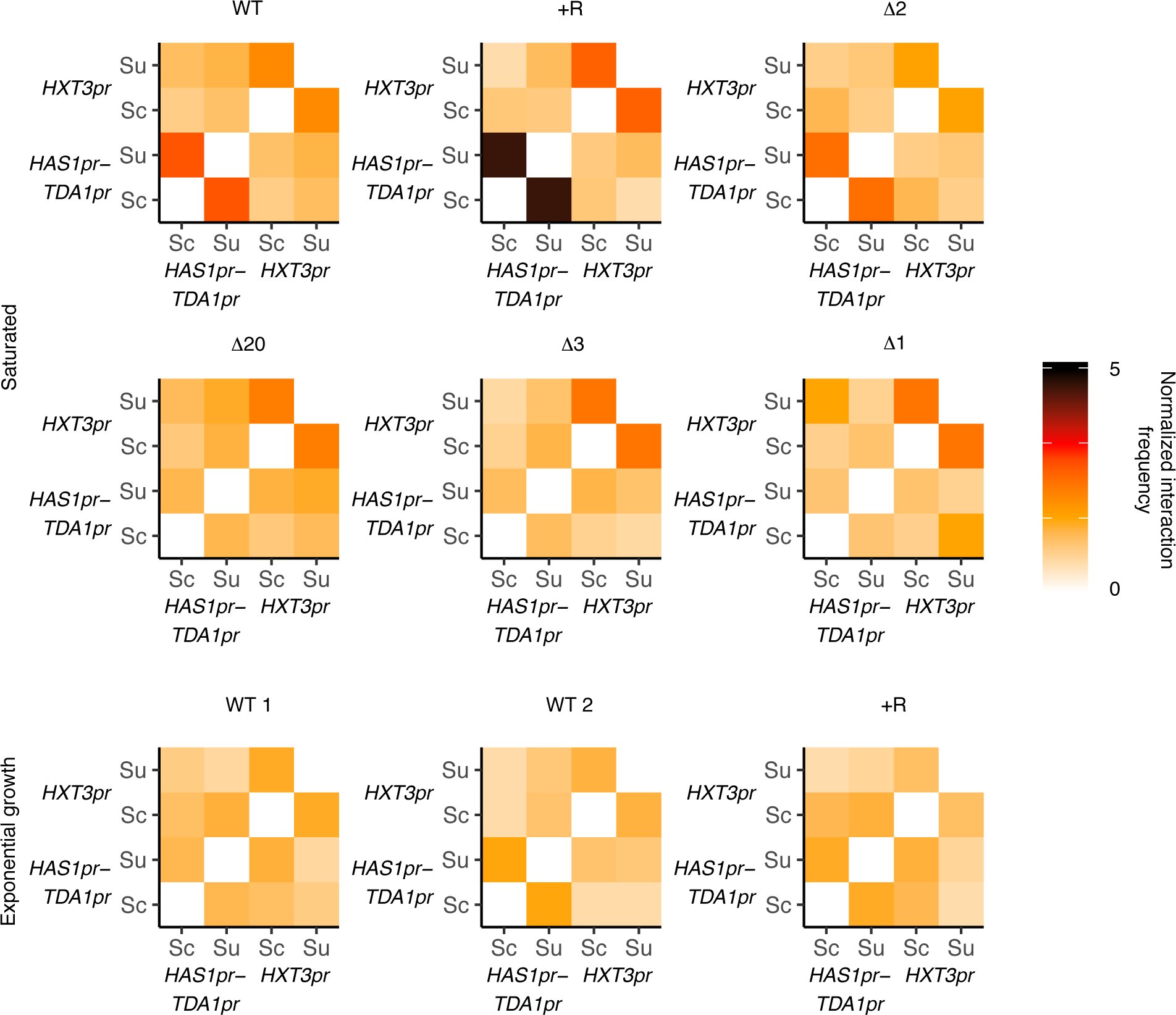
***HAS1pr-TDA1pr* and *HXT3pr* pairing are independent.** Hi-C interaction frequencies among the *S. cerevisiae* and *S. uvarum* copies of *HAS1pr-TDA1pr* and *HXT3pr*, at 32 kb resolution, across strains in pairing (saturated) and non-pairing (exponential growth) conditions. Sc indicates *S. cerevisiae*, and Su indicates *S. uvarum*. WT represents strain ILY456. +R indicates a restriction site added upstream of *HAS1* (YMD3920), ∆20 indicates a 20 kb deletion centered at *S. cerevisiae HAS1* (YMD3266), ∆3 indicates a 3 kb deletion centered at *S. cerevisiae HAS1* (YMD3267), ∆2 indicates a 2 kb deletion of the *S. cerevisiae HAS1* coding sequence (YMD3268), and ∆1 indicates a 1 kb deletion of the *S. cerevisiae HAS1pr-TDA1pr* intergenic region (YMD3269).

